# Glucosamine-6P and glucosamine-1P, respectively an activator and a substrate of rhodococcal ADP-glucose pyrophosphorylases, show a hint to ascertain (actino)bacterial glucosamine metabolism

**DOI:** 10.1101/2020.04.11.018549

**Authors:** AE Cereijo, HM Alvarez, AA Iglesias, MD Asencion Diez

**Affiliations:** Instituto de Agrobiotecnología del Litoral (UNL-CONICET), Facultad de Bioquímica y Ciencias Biológicas, CCT-Santa Fe, Colectora Ruta Nac 168 km 0, 3000 Santa Fe, Argentina; Instituto de Biociencias de la Patagonia (INBIOP), Universidad Nacional de la Patagonia San Juan Bosco y CONICET, Km 4-Ciudad Universitaria 9000, Comodoro Rivadavia, Chubut, Argentina

**Author notes:** Corresponding Author: Matias D. Asencion Diez, Instituto de Agrobiotecnología del Litoral (UNl-CONICET). CCT-Santa Fe, Colectora Ruta Nac 168 km 0, Santa Fe (3000), Argentina.

## Abstract

*Rhodococcus* spp. are important microorganisms for biotechnological purposes, such as bioremediation and biofuel production. The latter, founded on the oleaginous characteristic (high lipid accumulation) exhibited by many *Rhodococcus* species when grown in certain carbon sources under low nitrogen availability. These bacteria accumulate glycogen during exponential growth, and the glucan plays a role as an intermediary metabolite for temporary carbon storage related to lipid metabolism. The kinetic and regulatory properties of the ADP-glucose pyrophosphorylase (ADP-GlcPPase) from *Rhodococcus jostii* supports this hypothesis. The enzyme was found able to use glucosamine-1P as an alternative substrate. Curiously, the activity with glucosamine-1P was sensitive to glucose-6P, the main activator of actinobacterial ADP-GlcPPases. Herein, we report the study of glucosamine-1P related to the activity and regulation of ADP-GlcPPases from *R. jostii* and *R. fascians*, with the finding that glucosamine-6P is also a significant activator. Glucosamine-6P, belonging to a node between carbon and nitrogen metabolism, was identified as a main regulator in Actinobacteria. Thus, its effect on rhodococcal ADP-GlcPPases reinforces the function proposed for glycogen as temporary carbon storage. Results indicate that the activity of the studied enzymes using glucosamine-1P as a substrate responds to the activation by several metabolites that improve their catalytic performance, which strongly suggest metabolic feasibility. Then, studying the allosteric regulation exerted on an alternative activity would open two scenarios for consideration: (i) the existence of new molecules/metabolites yet undescribed, and (ii) evolutionary mechanisms underlying enzyme promiscuity that give rise new metabolic features in bacteria.

## Introduction

The accumulation of storage compounds allows to members of the *Rhodococcus* genus survival during changing environmental scenarios, involving high capacity for metabolic reconfiguration to adaptation [1,2]. The buildup of triacylglycerols is a common feature among actinobacteria belonging to the *Rhodococcus* genus. For example, *R. jostii* RHA1, produces high amounts (up to 60%) of lipids under nutrient starvation, thus being an oleaginous bacterium [1–3]. The latter represents a relevant biotechnological tool and positions rhodococci as critical for bioconversion processes to obtain biofuels precursors from simple carbon sources [1,4]. On the other hand, *R. fascians* lacks such oleaginous behavior, unless when grown in glycerol, after which is suitable for recycling of by-product waste from biodiesel production [4,5].

Glycogen synthesis was evidenced in several rhodococcal species [3]. The fact that the glucan accumulates during the exponential phase of growth supports its function as a temporary reserve of carbon that is mobilized in later stages. In other words, the glucan is an “intermediate metabolite” rather than a long-term storage compound, given that its accumulation is lower to 5% (w/w). Indeed, when lipid/fatty acids biosynthesis is disrupted in *R. opacus* - an oleaginous species like *R. jostii*-, carbon overflows toward its accumulation as glycogen [3]. Besides, the biochemical characterization of the enzyme catalyzing the key step in glycogen biosynthesis in *R. jostii* sustains this hypothesis (see below, and [6]).

Glycogen metabolism in bacteria depends on the availability of ADP-glucose (ADP-Glc), the specific glucosyl donor in its elongation. The sugar nucleotide is produced by ADP-Glc pyrophosphorylase (EC 2.7.7.27, ADP-GlcPPase), the rate-limiting enzyme in the glucan biosynthesis [7,8]. Except for the case of *Bacillus* spp., ADP-GlcPPases from all sources characterized so far are allosteric enzymes regulated by key metabolites from the central carbon pathway(s) in the respective organism [7,8]. Depending on the metabolite behaving as a regulator, ADP-GlcPPases were separated into nine groups [7], although such a classification is incomplete since it lacked data from Gram-positive organisms. We recently fill that gap by characterizing the enzyme from Actinobacteria (Gram-positive with high GC content DNA) and Firmicutes, finding distinctive allosteric effectors [6,9–12], which could add groups to that classification. Particularly, we established that glucose-6P (Glc-6P) is the primary activator in the ADP-GlcPPase from Actinobacteria and NADPH is a crucial inhibitor, thus connecting fatty acids synthesis with carbon to be allocated as glycogen [6,11]. The ADP-GlcPPases from *Streptomyces coelicolor* and *R. jostii* presented a broad sensitivity towards allosteric effectors according to organisms with broad metabolic capacities. There, Glc-6P stands as a “leading activator”, exerting the highest activation (*S. coelicolor*) [11] or exhibiting the highest affinity (*R. jostii*) [6]. When the ADP-GlcPPase from *R. jostii* (*Rjo*ADP-GlcPPase) was analyzed regarding its ability to use alternative sugar-1Ps, glucosamine-1P (GlcN-1P) was revealed as a substrate and, remarkably, was the only (besides Glc-1P) activity being activated by Glc-6P [6].

The above described particular characteristics of ADP-GlcPPase from rhodococci leads us to reconsider the relevance of GlcN metabolism and the role that GlcN-1P may play in these bacteria. Perhaps, such a function is different than a mere intermediary between the primary glycolytic pathways in the organism and the synthesis on UDP-acetylglucosamine (UDP-GlcNAc). In this work, we show that GlcN-1P is an efficient substrate for the ADP-GlcPPases from *R. jostii* and *R. fascians*, being this activity highly increased in the presence of GlcN-6P, a central metabolite in Actinobacteria [13].

## Results and Discussion

We previously reported that *Rjo*ADP-GlcPPase distinctively uses GlcN-1P as a substrate [6]. Although such activity is one order of magnitude lower respect to the use of Glc-1P, it is relevant that for both substrates, Glc-6P exerts a significant allosteric activation of the enzyme activity. Further analysis of these characteristics, driven to explore if whether or not the activation of the GlcN-1P-dependent activity of the rhodococcal enzyme is limited to Glc-6P, would be particularly useful. Thus, we assayed *Rjo*ADP-GlcPPase activity with 1 mM of Glc-1P or GlcN-1P in the presence of a single concentration (1 mM) of metabolites characterized as activators of the orthologous enzyme from different organisms [6–8,10]. We confirmed that all activators of the canonical reaction using Glc-1P also increased the activity with the alternative substrate GlcN-1P (Figure 1). Glc-6P, GlcN-6P, and Man-6P elicited the highest activation effects (more than 10-fold), whereas Fru-6P and phospho*enol*pyruvate (PEP) activated by about 4- and 8-fold, respectively. All effectors but PEP showed higher relative activation for GlcN-1P than for assays with Glc-1P.

**Figure 1:**
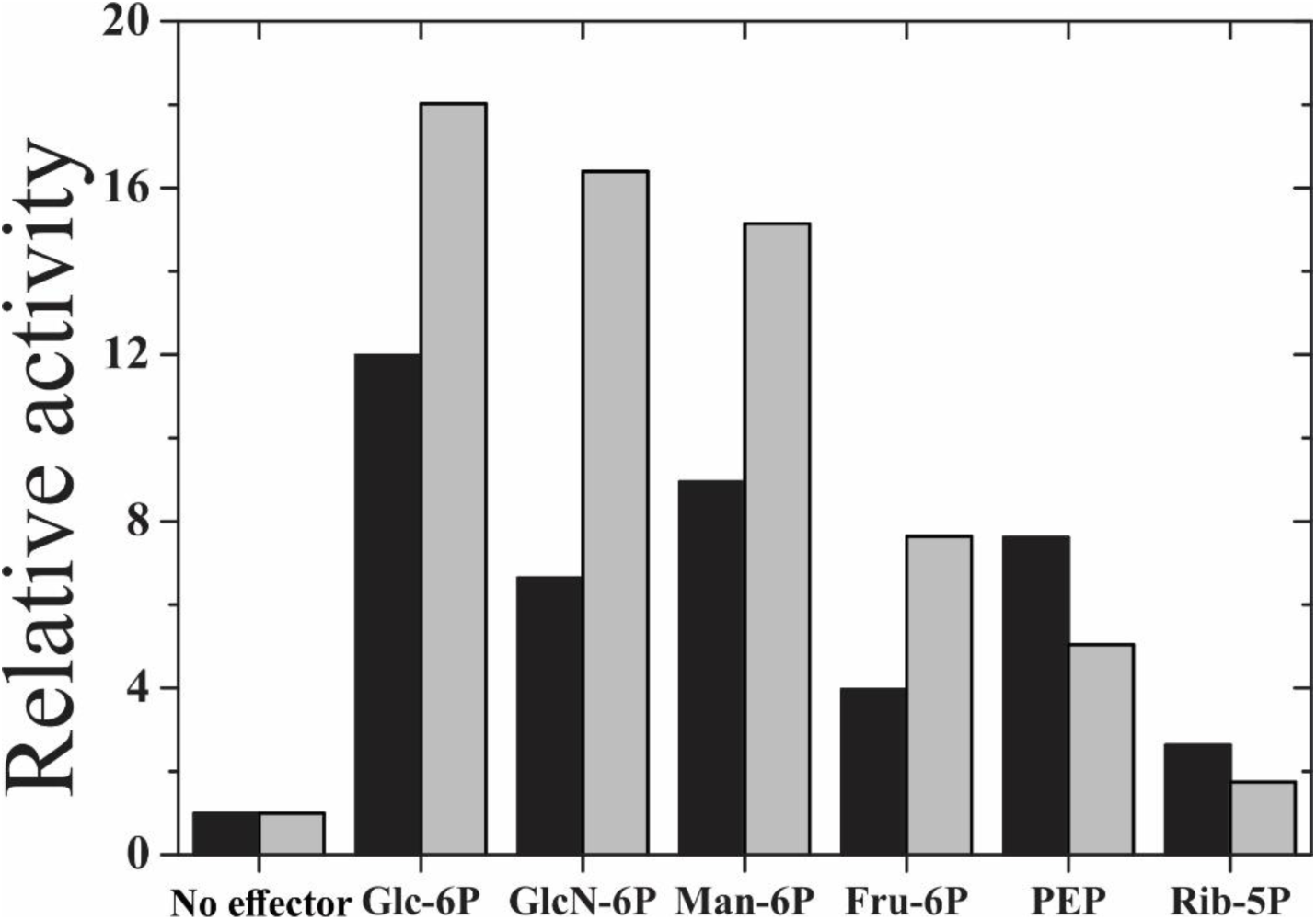
Relative activity of *Rjo*ADP-GlcPPase using Glc-1P (black bars) or GlcN-1P (grey bars) as a substrate in the presence of 1 mM of each effector. The activity in the absence of effector was 0.39 U/mg or 0.023 U/mg with Glc-1P or GlcN-1P, respectively.

Our results assign a critical role of GlcN-6P and GlcN-1P in the metabolism of *R. jostii*. On this basis, we performed a more detailed study of these amino sugars as allosteric effector and substrate of *Rjo*ADP-GlcPPase and made a comparison with the effects of Glc-6P and Glc-1P. Table 1 details the kinetic parameters determined for the activation of the enzyme by Glc-6P and GlcN-6P. As shown, *RjoA*DP-GlcPPase exhibited similar *A*_0.5_ values for each regulator regardless of whether the substrate is Glc-1P or GlcN-1P. In this context, the relative affinity of the enzyme for Glc-6P was one order of magnitude higher compared to GlcN-6P. Besides, both activators enhanced by ∼10-fold the acknowledged activity with Glc-1P and by ∼20-fold that with the alternative substrate GlcN-1P. Noteworthy, the GlcN-1P-dependent activity in the absence of activator is significantly low, depicting kinetics with incomplete saturation for substrates and then, the positive effect of the allosteric activation on the catalysis becomes more relevant. From these results, it is tempting to speculate that GlcN-6P would represent a signal of high carbon/energy availability in rhodococci, as proposed for other ADP-GlcPPase activators in many bacteria and plants [7,8]. The hypothesis also agrees with hexose-6P (and its amino sugar related) representing a critical metabolic node and itself an allosteric regulator of the activity of actinobacterial ADP-GlcPPase [9,11], particularly in *R. jostii* [6].

**Table 1:** Kinetic parameters for the activation of rhodococcal ADP-GlcPPases activity with Glc-1P or GlcN-1P as a substrate, and activated by Glc-6P or GlcN-6P

| Organism | Effector | Glc-1P | GlcN-1P | Glc-1P | GlcN-1P | Glc-1P | GlcN-1P |
| --- | --- | --- | --- | --- | --- | --- | --- |
| | | $A_{0.5}$ (mM) | | $k_{cat}$ (s<sup>-1</sup>) | | Activation (-fold) | |
| *R. jostii* | Glc-6P | 0.088 ± 0.005 | 0.059 ± 0.001 | 3.41 | 0.46 | 12 | 18 |
|  | GlcN-6P | 0.57 ± 0.05 | 0.56 ± 0.02 | 2.52 | 0.51 | 9 | 20 |
| *R. fascians* | Glc-6P | 0.12 ± 0.01 | 0.018 ± 0.003 | 22.9 | 0.33 | 2.6 | 12.8 |
|  | GlcN-6P | 0.42 ± 0.03 | 0.054 ± 0.009 | 20.2 | 0.29 | 2.3 | 11.6 |

We previously demonstrated that the allosteric effectors of *Rjo*ADP-GlcPPase increased its affinity towards substrates, remarkably Glc-1P [6]. In the present study GlcN-6P behaved similarly since it (at 1.5 mM in the assay media) decreased the *S*_0.5_ for Glc-1P from 1.8 mM to 0.073 mM (Figure 2A), This result, combined with activation fold detailed in Table 1, indicates that GlcN-6P increases the *Rjo*ADP-GlcPPase catalytic efficiency (measured as *k*_cat_/*S*_0.5_, see [6,7,10]) by more than two orders of magnitude. The latter supports a link between glycogen and amino sugars metabolism, where an increase in GlcN-6P (either by exogenous intake of GlcN or by peptidoglycan degradation [13]) might trigger the temporary accumulation of carbon in the glycogen form. This scenario agrees with the metabolic role proposed for the polyglucan in this oleaginous bacterium [2,3]. Besides, we established that GlcN-6P also activates the *Rjo*ADP-GlcPPase catalysis for the use of GlcN-1P (Figure 2B), sustaining the idea of an effector/substrate relationship similar to the canonical Glc-6P/Glc-1P. Certainly, GlcN-6P was an activator with a relevant effect on the GlcN-1P catalysis, allowing the enzyme to reach a *k*_cat_ comparable to that with Glc-6P (Table 1). Besides, such a turnover number supports the feasibility of the reaction with GlcN-1P in the bacterium cell [14]. Thus, we propose that *Rjo*ADP-GlcPPase would adapt to a scenario where the GlcN availability activates the drive of amino sugar-1P toward the synthesis of ADP-GlcN, which would further follow a yet unknown metabolic fate.

**Figure 2:**
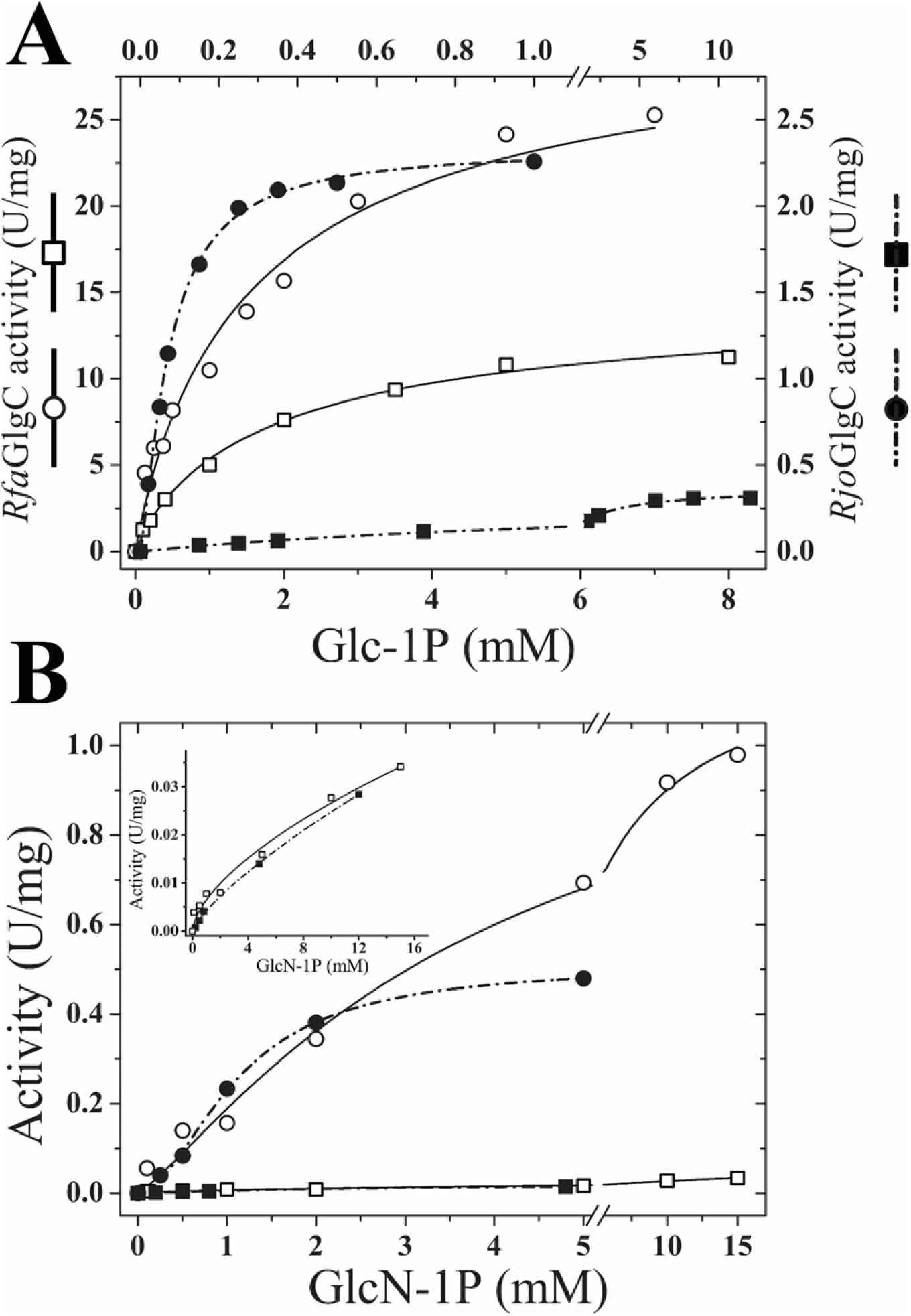
Saturation curves of Glc-1P (A) consumption by *Rjo*ADP-GlcPPase (right axis – filled symbols) and *Rfa*ADP-GlcPPase (left axis – open symbols) in absence (squares) or presence of 1 mM GlcN-6P (circles) and 1 mM ATP. GlcN-1P curves (B) for both *Rjo*ADP-GlcPPase (filled symbols) and *Rfa*ADP-GlcPPase (open symbols) were assayed without effector (squares) or in presence of 1 mM GlcN-6P (circles).

We extended the study to determine the kinetic and regulatory properties of the ADP-GlcPPase from *R. fascians* (*Rfa*ADP-GlcPPase) which is a close related organism with different metabolic behavior, mainly regarding lipid accumulation [1]. Performing this type of characterization in the ADP-GlcPPase from close organisms in the *Rhodococcus* genus would be critical to understand the role of GlcN-1P (as a substrate) and GlcN-6P (as an allosteric effector) in carbohydrate metabolism. The *Rfa*ADP-GlcPPase was recombinantly produced with the same strategy used for *Rjo*ADP-GlcPPase [6]. Its kinetic parameters were determined, depicting almost hyperbolic saturation curves for substrates with *S*_0.5_ values of 1.17 and 0.58 mM for Glc-1P and ATP, respectively (Table 2). Remarkably, the enzyme showed a *V*_max_ (12 U/mg) 30-fold higher compared to *Rjo*ADP-GlcPPase [6], thus being the most active actinobacterial ADP-GlcPPase characterized so far. As well, it was activated by Glc-6P, Man-6P, Fru-6P and PEP (data not shown), in a similar way to that determined for *Rjo*ADP-GlcPPase. A minor difference is that the activation effect is lower for the *R. fascians* enzyme. Indeed, all effectors increased the enzyme *V*_max_ between 2- and 3-fold, resembling the activation of the mycobacterial ADP-GlcPPase [9]. GlcN-6P also activated the *R. fascians* enzyme (Figure 2). We determined that Glc-6P and GlcN-6P respectively enhanced by ∼6-fold and ∼2-fold the catalytic efficiency of the enzyme (Table 2).

**Table 2:** Kinetic parameters for Glc(N)-1P utilization by *Rf*aADP-GlcPPase in absence or presence of Glc(N)-6P effectors. *nd\**: not determined, since no saturation curves were obtained (see Figure 2B).

| Substrate | Effector | $S_{0.5}$ (mM) | $k_{cat}$ (s<sup>-1</sup>) | $k_{cat}/S_{0.5}$ (s<sup>-1</sup> mM<sup>-1</sup>) |
| --- | --- | --- | --- | --- |
| **Glc-1P** | None | $1.17 \pm 0.08$ | 8.8 | 7.5 |
| | Glc-6P (1 mM) | $0.59 \pm 0.04$ | 24.3 | 41.1 |
| | GlcN-6P (1 mM) | $1.41 \pm 0.12$ | 21.3 | 15.2 |
| **GlcN-1P** | None | *nd\** | *nd\** | *nd\** |
| | Glc-6P (1 mM) | $3.4 \pm 0.5$ | 2.11 | 0.62 |
| | GlcN-6P (1 mM) | $3.7 \pm 0.6$ | 0.87 | 0.23 |

Figure 2B shows that *Rfa*ADP-GlcPPase is also able to use GlcN-1P as a substrate, exhibiting non-saturation kinetics, as observed for the *Rjo*ADP-GlcPPase. Similar specific activity values (∼30 mU/mg) were achieved for both rhodococcal ADP-GlcPPases at the highest GlcN-1P concentration assayed. As shown in Table 1, the GlcN-1P dependent activity of *Rfa*ADP-GlcPPase was enhanced by Glc-6P and GlcN-6P (see also Figure 2). The kinetic parameters of the activation with each of the substrates detailed in Table 1 indicate that Glc-6P and GlcN-6P behave similarly in the regulation of the *R. fascians* enzyme. Thus, for both activators, the *A*_0.5_ was lower and the activation fold higher when the substrate was GlcN-1P, compared to Glc-1P. These results reinforce the view that GlcN-6P constitutes an efficient activator of *Rfa*ADP-GlcPPase, and it would signal a high carbon availability to be led to glycogen synthesis, as we above described for the *Rjo*ADP-GlcPPase. From data in Table 1, it is worthy to calculate the specificity constant for each activator, defined as the ratio between net activation and the respective *A*_0.5_ [15], which evaluates the capacity of the respective allosteric effector with the different substrates. Figure 3 illustrates that for *Rjo*ADP-GlcPPase, the Glc-6P specificity constant is higher (∼9-fold) compared to GlcN-6P with either substrate. For both activators, the parameter is ∼2-fold higher regarding the GlcN-1P activation than for Glc-1P. In *Rfa*ADP-GlcPPase, the ratio between the specificity constants of Glc-6P over GlcN-6P is in the range 3 to 4; however (and interestingly), both activator parameters are more than one order of magnitude higher for the use of GlcN-1P than to Glc-1P (note the log scale in Figure 3). These results strongly suggest that in rhodococci, the allosteric regulation of ADP-GlcPPase by Glc-6P and GlcN-6P enhance the canonical activity with Glc-1P and in a higher degree than with the alternative amino sugar substrate. The described property regarding the allosteric regulation of GlcN-1P activity is more significant in certain species of *Rhodococcus*, like *R. fascians*, that exhibits critically distinctive metabolic capacities.

**Figure 3:**
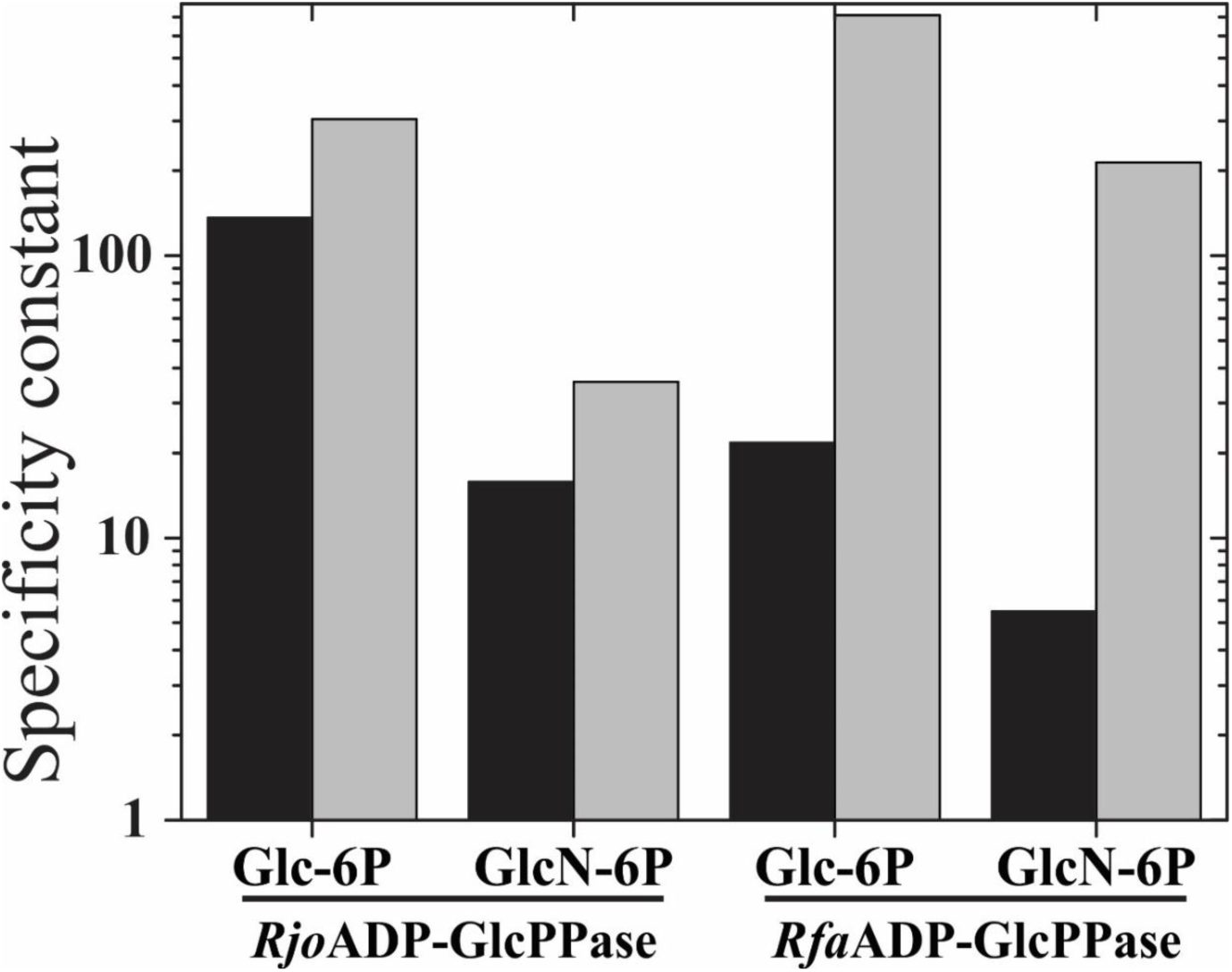
Histogram presenting the specificity constant for both enzymes determined for Glc-1P (black bars) or GlcN-1P (grey bars) in presence of the effectors Glc-6P or GlcN-6P.

The ability of ADP-GlcPPases from *R. jostii* and *R. fascians* in using GlcN-1P as a substrate, and this activity being sensitive to allosteric regulation, sustains the occurrence of a particular metabolic context inside the bacteria cell [14]. This hypothesis is also supported by the proposal that turnover numbers (*k*_cat_) determined *in vitro* are quite similar to those taking place *in vivo* [16]. In this context, the ADP-GlcN synthesis inside rhodococci cells could be inferred, although this framework needs of future experimental studies (actually, ongoing experiments in our group). Previously, we demonstrated that a new type of nucleotide-sugar pyrophosphorylase (GDP-GlcPPase, EC 2.7.7.34) from *Streptomyces venezuelae* (a close related organism to rhodococci) produced metabolically significant amounts of the specific glucosyl donor (GDP-Glc) for trehalose synthesis [17]. This new enzyme showed an efficiency toward GDP-Glc synthesis in the order of 10^−1^ s^-1^ mM^-1^, similar to parameters presented in this work for both rhodococcal ADP-GlcPPases sustaining our proposed scenario. Consequently, it appears acceptable to assume that these two enzymes are proficient at generating an NDP-sugar molecule, ADP-GlcN, never described so far. Further support is given by the fact that Glc-6P and GlcN-6P (the latter reported as critical in actinobacterial metabolism [13]), strongly activate GlcN-1P catalysis by increasing the catalytic efficiency for this substrate and/or by boosting the specificity constant for the effector (notoriously mainly for *Rfa*ADP-GlcPPase).

Results presented *ut supra* would have implications at two levels. First, either *Rjo*ADP-GlcPPase or *Rfa*ADP-GlcPPase could constitute a biotechnological tool for *in vitro* synthesis and purification of novel ADP-(amino)sugars, which could be used as substrates to explore for (amino)glucosyl-transferase activities [18] or applied to *glycorandomization* processes [19,20]. Last but not least, the consideration in the *Rhodococcus* genus of alternative (new) metabolic pathway(s), possibly driving high carbon availability (signaled by GlcN-6P) to a yet non-established intracellular fate. Even more, a novel metabolic node could be proposed at the GlcN-1P level. This metabolite, rather than being a simple intermediary between Fru-6P and UDP-GlcNAc interconversion, could be the starting point of pathway(s) to harness the carbon available by GlcN intake and/or peptidoglycan consumption, by storing it as glycogen (or a similar glucan) or a new carbohydrate-related molecule. Also, the example presented here could be a case-study of underground metabolism [21–23], where the promiscuity of an ADP-GlcPPase evolved a secondary activity with an allosteric sensitivity to effectors, thus allowing an *a priori* non-relevant metabolite to be part of a physiological reactions and a metabolic node. Finally, these rhodococcal ADP-GlcPPases constitute an attractive model to study the evolutionary mechanisms involved in promiscuity to allosterism relationships.

## Remarks

We show that GlcN-1P and GlcN-6P are (respectively) an efficient substrate and allosteric activator of two rhodococcal ADP-GlcPPases. The *Rhodococcus* species exhibit differential metabolic behaviors, sustaining the idea that the glucan plays a temporary carbon storage role, as hypothesized before [2,3,6]. Indeed, GlcN-6P is at the corner of carbon and nitrogen metabolism, as well as the cell wall synthesis via UDP-GlcNAc [13], thus reinforcing the critical role that glycogen has as an intermediary molecule. Also, a close link between amino sugar availability and glycogen metabolism (never reported so far) could be postulated. This fact would be a signal for carbon storage, later to be directed to lipid biosynthesis. A clear similitude could be deduced between Glc-6P/Glc-1P and GlcN-6P/GlcN-1P, probably also involving the same evolutionary mechanism.

Remarkably, GlcN-1P arises as an efficient substrate for rhodococcal ADP-GlcPPases, where the kinetic parameters sustain intracellular feasibility towards ADP-GlcN synthesis. GlcN-1P consumption could be the starting point of new metabolic pathways and also, a new carbon node for yet unknown metabolic fates. Still, questions remain to be answered, such as, is ADP-GlcN synthesized *in vivo*? Which is its intracellular fate? Could be an aminated glycogen synthesized in these rhodococcal cells? Solving these issues would be critical to further understand the complex metabolism presented by *Rhodococcus* genus and to improve their application as biotechnological tools for bioremediation and/or bioconversions processes. Thus, the present report opens the door for further biochemical and metabolic exploration in Actinobacteria.

## Acknowledgments

This work was supported by grants from ANPCyT (PICT’17 1515 to AAI and PICT’15 0634 to MDAD) and CONICET (PIP2015-2016 0529 to HMA and PUE 2016 238 to IAL). AEC is a Fellow from CONICET. HMA, AAI and MDAD are Career Investigator members from the same Institution

## Methods

Protein standards, antibiotics, IPTG, Glc-1P, Glc-6P, GlcN-1P, ATP, Fru-6P, PEP and oligonucleotides were obtained from Sigma-Aldrich (St. Louis, MO, USA). All other reagents were of the highest quality available.

*Escherichia coli* Top 10 F′ cells (Invitrogen) and pGEM^®^T Easy vector (Promega) were used for cloning procedures. The *glgC* gene from *R. fascians* (*RfaglgC*) was expressed in *E. coli* BL21 (DE3; Invitrogen) using pET28c vector (Novagen). DNA manipulations, *E. coli* cultures as well as transformations were performed according to standard protocols [24].

The *glgC* gene (GeneBank: KMJ51055.1) coding for ADP-Glc PPase from *R. fascians* was amplified by PCR using genomic DNA as template. Primers were designed according to available genomic information (BioProject ID 286803) in the GenBank database. The forward primer (5′-<u>CATATG</u>AGGACACAACCGCACGTAC-3′) introduced a *Nde*I site (underlined) using the initiation codon (changing GTG to ATG). The reverse primer introduced a *Hind*III site (underlined) downstream a stop codon: 5′-<u>AAGCTT</u>CTAGATCCAGACGCCCTTAC-3′. PCR reaction mixtures (50 μl) contained 100 ng of genomic DNA; 0.2 mM of each dNTP; 5 mM Mg2+, 50 pmol of each primer and 1 U *Pfu* DNA polymerase (Fermentas). Standard conditions of PCR were used for 30 cycles: denaturation at 94°C for 1 min; annealing at 56°C for 30 s, and extension at 72°C for 2 min, with a final extension of 10 min at 72°C. PCR reaction mixtures were solved in 1% (w/v) agarose gel and purified by means of Wizard SV gel and PCR Clean Up kits (Promega). The amplified gene was treated with *Taq* polymerase (Fermentas) and then cloned into pGEM-T Easy. The identity of the cloned gene was determined by full sequencing. (Macrogen, South Korea). Afterward, the pGEM-T Easy plasmid harboring *RfaglgC* coding sequence was digested with *Nde*I and *Hind*III and the released gene was cloned into pET28c.

Competent *E. coli* BL21 (DE3) cells were transformed with the [pET28c/*RfaglgC*] construction. Protein expression was carried out using LB medium (10 g/l tryptone; 5 g/l yeast extract; 5 g/l NaCl) supplemented with 100 g/ml kanamycin. Cells grown at 37°C and 250 rpm until OD_600_ ∼0.6 were induced for 16 h at 18°C after IPTG addition (0.2 mM). Then, cells were harvested by centrifugation at 6000 ×*g* for 5 min and stored at −20°C until use.

Purification techniques were done at 4°C. Cells were resuspended in *Buffer H* [25 mM Tris-HCl pH 8.0, 300 mM NaCl, 10 mM imidazole] and after sonication (on ice. 5 s pulse on with intervals of 3 s pulse off) for of 10 min, the resulted suspension was centrifuged twice at 30000 ×*g* for 15 min. The supernatant was loaded onto a 1 ml HisTrap column (GE Healthcare) previously equilibrated with *Buffer H*. The recombinant *Rfa*ADP-GlcPPase eluted with a linear gradient from 10 to 300 mM imidazole in *Buffer H* (40 volumes). Fractions containing the highest activity were pooled and concentrated to 1-2 ml. Active *Rfa*ADP-GlcPPase fractions were dialyzed against *Buffer X* [50 mM HEPES pH 8.0, 0.1 mM EDTA, 0.5 mM DTT, 20% (w/v) sucrose], flash-freeze and stored at −80°C until use, being fully actives for 3 months. In addition, *Rjo*ADP-GlcPPase was obtained as described before [6]. Protein concentration was determined by a modified Bradford assay [25]. Purified and purification fractions of *Rfa*ADP-GlcPPase were defined by sodium dodecyl sulfate polyacrylamide gel electrophoresis (SDS-PAGE) according to [26].

Activity was determined at 37°C in the ADP-Glc synthesis direction by following the formation of P_i_ (after hydrolysis of PP_i_ by inorganic pyrophosphatase) by a colorimetric method [27]. Reaction mixtures contained 50 mM MOPS pH 8.0, 10 mM MgCl_2_, 1.5 −2 mM ATP, 0.2 mg/ml bovine serum albumin, 0.5 U/ml yeast inorganic pyrophosphatase and a proper enzyme dilution. Assays were initiated by addition of Glc-1P or GlcN-1P in a total volume of 50 μl. The reaction mixtures were incubated at 37°C for 10 min and terminated by adding the Malachite Green reactive. The complex formed with the released P_i_ was measured at 620 nm. One unit of activity (U) is defined as the amount of enzyme catalyzing the formation of 1 μmol of product per min, under conditions above described in each case.

Saturation curves were obtained after assaying enzyme activity at different concentrations of the variable substrate or effector and saturating levels of the others. Experimental data were plotted as enzyme activity (U/mg) *versus* substrate (or effector) concentration (mM), and kinetic constants were determined by fitting the data to the Hill equation as described elsewhere [28]. We analyzed enzyme behavior by fitting with the Levenberg–Marquardt non-linear least-squares algorithm provided by the computer program Origin™. Hill plots were used to calculate the Hill coefficient (*n*_H_), the maximal velocity (*V*_max_), and the kinetic constants that correspond to the activator or substrate, concentrations giving 50% of the maximal activation (*A*_0.5_) or velocity (*S*_0.5_), respectively. The specificity constant for activators were obtained as reported in [15], defined as the ratio between net activation the respective *A*_0.5_. The net activation was calculated as (*V*_max_-*v*_0_)/*v*_0_, where *V*_max_ is the activity obtained in presence of the respective effector while *v*_0_ is the activity without effector. The specificity constant parameter is analogous to the catalytic efficiency, used to compare the substrate specificity of enzymes with simple hyperbolic kinetics. Parameters are the mean of three independent sets of data reproducible within ±10%.

## References

[1] H.M. Alvarez, O.M. Herrero, R.A. Silva, M.A. Hernández, M.P. Lanfranconi, M.S. Villalba, Insights into the Metabolism of Oleaginous *Rhodococcus* spp, Appl. Environ. Microbiol., 85 (2019).

[2] M.A. Hernández, W.W. Mohn, E. Martínez, E. Rost, A.F. Alvarez, H.M. Alvarez, Biosynthesis of storage compounds by *Rhodococcus jostii* RHA1 and global identification of genes involved in their metabolism, BMC Genomics, 9 (2008) 600.

[3] M.A. Hernandez, H.M. Alvarez, Glycogen formation by *Rhodococcus* species and the effect of inhibition of lipid biosynthesis on glycogen accumulation in *Rhodococcus opacus* PD630, FEMS Microbiol. Lett., 312 (2010) 93–99.

[4] O.M. Herrero, M.S. Villalba, M.P. Lanfranconi, H.M. Alvarez, *Rhodococcus* bacteria as a promising source of oils from olive mill wastes, World J. Microbiol. Biotechnol., 34 (2018) 114.

[5] O.M. Herrero, G. Moncalián, H.M. Alvarez, Physiological and genetic differences amongst *Rhodococcus* species for using glycerol as a source for growth and triacylglycerol production, Microbiology, 162 (2016) 384–397.

[6] A.E. Cereijo, M.D. Asencion Diez, J.S. Davila Costa, H.M. Alvarez, A.A. Iglesias, On the Kinetic and Allosteric Regulatory Properties of the ADP-Glucose Pyrophosphorylase from *Rhodococcus jostii:* An Approach to Evaluate Glycogen Metabolism in Oleaginous Bacteria, Front. Microbiol., 7 (2016) 830.

[7] M.A. Ballicora, A.A. Iglesias, J. Preiss, ADP-glucose pyrophosphorylase, a regulatory enzyme for bacterial glycogen synthesis, Microbiol. Mol. Biol. Rev., 67 (2003) 213–25, table of contents.

[8] M.A. Ballicora, A.A. Iglesias, J. Preiss, ADP-glucose pyrophosphorylase: A regulatory enzyme for plant starch synthesis, Photosynth. Res., 79 (2004) 1–24.

[9] M.D. Asención Diez, A.M. Demonte, K. Syson, D.G. Arias, A. Gorelik, S.A. Guerrero, S. Bornemann, A.A. Iglesias, Allosteric regulation of the partitioning of glucose-1-phosphate between glycogen and trehalose biosynthesis in *Mycobacterium tuberculosis*, Biochim. Biophys. Acta, 1850 (2015) 13–21.

[10] A.E. Cereijo, M.D. Asencion Diez, M.A. Ballicora, A.A. Iglesias, Regulatory Properties of the ADP-Glucose Pyrophosphorylase from the Clostridial Firmicutes Member *Ruminococcus albus*, J. Bacteriol., 200 (2018).

[11] M.D. Asencion Diez, S. Peiru, A.M. Demonte, H. Gramajo, A.A. Iglesias, Characterization of recombinant UDP- and ADP-glucose pyrophosphorylases and glycogen synthase to elucidate glucose-1-phosphate partitioning into oligo- and polysaccharides in *Streptomyces coelicolor*, J. Bacteriol., 194 (2012) 1485–1493.

[12] M.D. Asención Diez, A.M. Demonte, S.A. Guerrero, M.A. Ballicora, A.A. Iglesias, The ADP-glucose pyrophosphorylase from *Streptococcus mutans* provides evidence for the regulation of polysaccharide biosynthesis in Firmicutes, Mol. Microbiol., 90 (2013) 1011–1027.

[13] H.U. van der Heul, B.L. Bilyk, K.J. McDowall, R.F. Seipke, G.P. van Wezel, Regulation of antibiotic production in Actinobacteria: new perspectives from the post-genomic era, Nat. Prod. Rep., 35 (2018) 575–604.

[14] A. Bar-Even, E. Noor, Y. Savir, W. Liebermeister, D. Davidi, D.S. Tawfik, R. Milo, The moderately efficient enzyme: Evolutionary and physicochemical trends shaping enzyme parameters, Biochemistry, 50 (2011) 4402–4410.

[15] M.L. Kuhn, C.M. Figueroa, A.A. Iglesias, M.A. Ballicora, The ancestral activation promiscuity of ADP-glucose pyrophosphorylases from oxygenic photosynthetic organisms, BMC Evol. Biol., 13 (2013) 51.

[16] D. Davidia, E. Noorb, W. Liebermeisterc, A. Bar-Evend, A. Flamholze, K. Tummlerf, U. Barenholza, M. Goldenfelda, T. Shlomig, R. Miloa, Global characterization of i*n vivo* enzyme catalytic rates and their correspondence to in vitro *k*_cat_ measurements, Proc. Natl. Acad. Sci. U. S. A., 113 (2016) 3401–3406.

[17] M.D. Asención Diez, F. Miah, C.E.M. Stevenson, D.M. Lawson, A.A. Iglesias, S. Bornemann, The Production and Utilization of GDP-glucose in the Biosynthesis of Trehalose 6-Phosphate by *Streptomyces venezuelae*, J. Biol. Chem., 292 (2017) 945–954.

[18] S.H. Park, H.Y. Park, J.K. Sohng, H.C. Lee, K. Liou, Y.J. Yoon, B.G. Kim, Expanding substrate specificity of GT-B fold glycosyltransferase via domain swapping and high-throughput screening, Biotechnol. Bioeng., 102 (2009) 988–994.

[19] V.S. Bais, S. Batra, B. Prakash, Identification of two highly promiscuous thermostable sugar nucleotidylyltransferases for *glycorandomization*, FEBS J., 285 (2018) 2840–2855.

[20] V. Křen, T. Řezanka, Sweet antibiotics - The role of glycosidic residues in antibiotic and antitumor activity and their randomization, FEMS Microbiol. Rev., 32 (2008) 858–889.

[21] R. D’Ari, J. Casadesús, Underground metabolism, Bioessays, 20 (1998) 181–6.

[22] A. Peracchi, The Limits of Enzyme Specificity and the Evolution of Metabolism, Trends Biochem. Sci., 43 (2018) 984–996.

[23] J. Rosenberg, F.M. Commichau, Harnessing Underground Metabolism for Pathway Development, Trends Biotechnol., 37 (2019) 29–37.

[24] J. Sambrook, D. Russell, Molecular cloning: A laboratory Manual, Cold Spring Harbor Laboratory Press, New York, 2001.

[25] M.M. Bradford, A rapid and sensitive method for the quantitation of microgram quantities of protein utilizing the principle of protein-dye binding, Anal. Biochem., 72 (1976) 248–254.

[26] U. Laemmli, Cleavage of Structural Proteins during the Assembly of the Head of Bacteriophage T4, Nature, 227 (1970) 680–685.

[27] C. Fusari, A.M. Demonte, C.M. Figueroa, M. Aleanzi, A.A. Iglesias, A colorimetric method for the assay of ADP-glucose pyrophosphorylase, Anal. Biochem., 352 (2006) 145–147.

[28] M.A. Ballicora, E.D. Erben, T. Yazaki, A.L. Bertolo, A.M. Demonte, J.R. Schmidt, M. Aleanzi, C.M. Bejar, C.M. Figueroa, C.M. Fusari, A.A. Iglesias, J. Preiss, Identification of regions critically affecting kinetics and allosteric regulation of the *Escherichia coli* ADP-glucose pyrophosphorylase by modeling and pentapeptide-scanning mutagenesis, J. Bacteriol., 189 (2007) 5325–5333.

